# Toward identification of common DNA repair process in mutational signatures

**DOI:** 10.64898/2026.01.29.702643

**Authors:** Damian Wójtowicz, Marcin Wierzbiński, Jan Hoinka, Roded Sharan, Teresa M. Przytycka

## Abstract

Mutational signatures are characteristic patterns of mutation frequencies assumed to be generated by specific mutagenic processes. A growing catalog of mutational signatures exists, but tools to systematically infer relationships between them remain limited. A mutational signature can be viewed as the combined outcome of two processes: DNA damage and DNA repair. Since cancer therapies often target DNA repair, inferring DNA repair pathways is important for treatment design, even when the mutagenic process is unknown.

Here, we model the DNA repair step as a transformation, called RePrint, from damaged nucleotides to repair-related mutation patterns conditioned on the damage. We demonstrate that RePrint similarity is indicative of shared DNA repair mechanisms, enabling *guilt-by-association* prediction of DNA repair pathways. Using experimentally annotated signatures from environmental exposures and CRISPR gene knockouts as gold standards, we demonstrate that RePrint-based clustering consistently outperforms signature-based clustering across multiple evaluation metrics. We validate several guilt-by-association predictions with literature evidence, demonstrating RePrint’s ability to identify shared repair mechanisms even among signatures with divergent mutational profiles.

RePrint provides the first approach to systematically transfer DNA repair information between signatures, opening doors to understanding signatures of unknown origin and informing therapeutic strategies. An open-source implementation is available at https://github.com/wojtowicz-lab/RePrintPy

## 1. Introduction

Cancer genomes accumulate a large number of somatic mutations resulting from various endogenous and exogenous causes, including normal DNA damage and repair, cancer-related aberrations of the DNA maintenance machinery, and mutations triggered by carcinogenic exposures. Different mutagenic processes lead to distinct patterns of somatic mutations called mutational signatures. These mutational signatures can be thought of as fingerprints of the corresponding mutagenic processes that act on a given genome (Nik-Zainal et al., 2012; Alexandrov et al., 2013a,b). Analysis of mutational signatures has emerged as an important approach to study the mutagenic processes that have shaped the evolution of a cancer genome and as a predictor of drug response (Kim et al., 2021; Van Hoeck et al., 2019).

Mutational signatures are defined based on a partition of mutations into mutation categories. Typically, it is assumed that somatic mutations fall into 96 categories corresponding to 6 types of mutation base substitutions and 16 combinations of flanking nucleotide residues (e.g., TCC>TTC represents the C>T mutation in the nucleotide context of T_C). The probability of generating a mutation of a given category depends on the mutational process. To reflect this, mutational signatures are defined as vectors of probability distributions over mutation categories. As the amount of cancer genome data increases, so does the set of signatures. Currently, the COSMIC dictionary of signatures contains about 100 such signatures. Many of these signatures have been linked to specific mutagenic processes with varying degrees of confidence and precision, ranging from mechanistic explanations to purely statistical associations between signatures and potential causes (Alexandrov et al., 2020). Identifying mutational signatures in new datasets and linking them with molecular, genomic, and environmental causes has become one of the cornerstones of computational analysis of cancer datasets (Kim et al., 2021).

Over the years, extensive knowledge has been accumulated about individual signatures; however, the tools available to systematically explore the relationships between them remain limited. The primary approach for leveraging existing knowledge to characterize a potentially novel signature is to test whether a newly inferred signature exhibits strong similarity (cosine or root-mean-square deviation) to the known signatures, thereby suggesting that the signature represents the same or a very similar mutagenic process. Additional features, such as strand asymmetry, which is manifested as mutational biases between leading and lagging strands, or transcribed and non-transcribed regions, may further indicate the involvement of specific DNA damage or repair mechanisms.Overall, the etiology of many of the signatures remains unknown, and it becomes crucial to define new relations that could be used to identify biological properties shared by signatures. Because cancer therapies often target DNA synthesis or repair, inferring DNA repair pathways active in a given mutational signature is of particular interest Van Hoeck et al. (2019).

In this paper, we introduce a new similarity measure between signatures, designed to identify mutagenic processes that are likely to share the DNA repair mechanism without the requirement that the DNA damage process be known. The measure is based on a transformation that converts mutational signatures into a vector of conditional probabilities, termed RePrint. Although RePrint has been introduced previously (Wojtowicz et al., 2020), its broader utility as a measure of similarity of DNA repair processes in different signatures was not explored. RePrint leverages the key conceptual framework that a mutational signature should be viewed as the combined outcome of two distinct processes: DNA damage and subsequent DNA repair (Wojtowicz et al., 2020, 2021; Volkova et al., 2020). Under this assumption, we can model the DNA repair step as a transformation from damaged nucleotides due to a mutagenic process to a characteristic mutation pattern conditioned on the damage (Figure 1).

**Figure 1.**
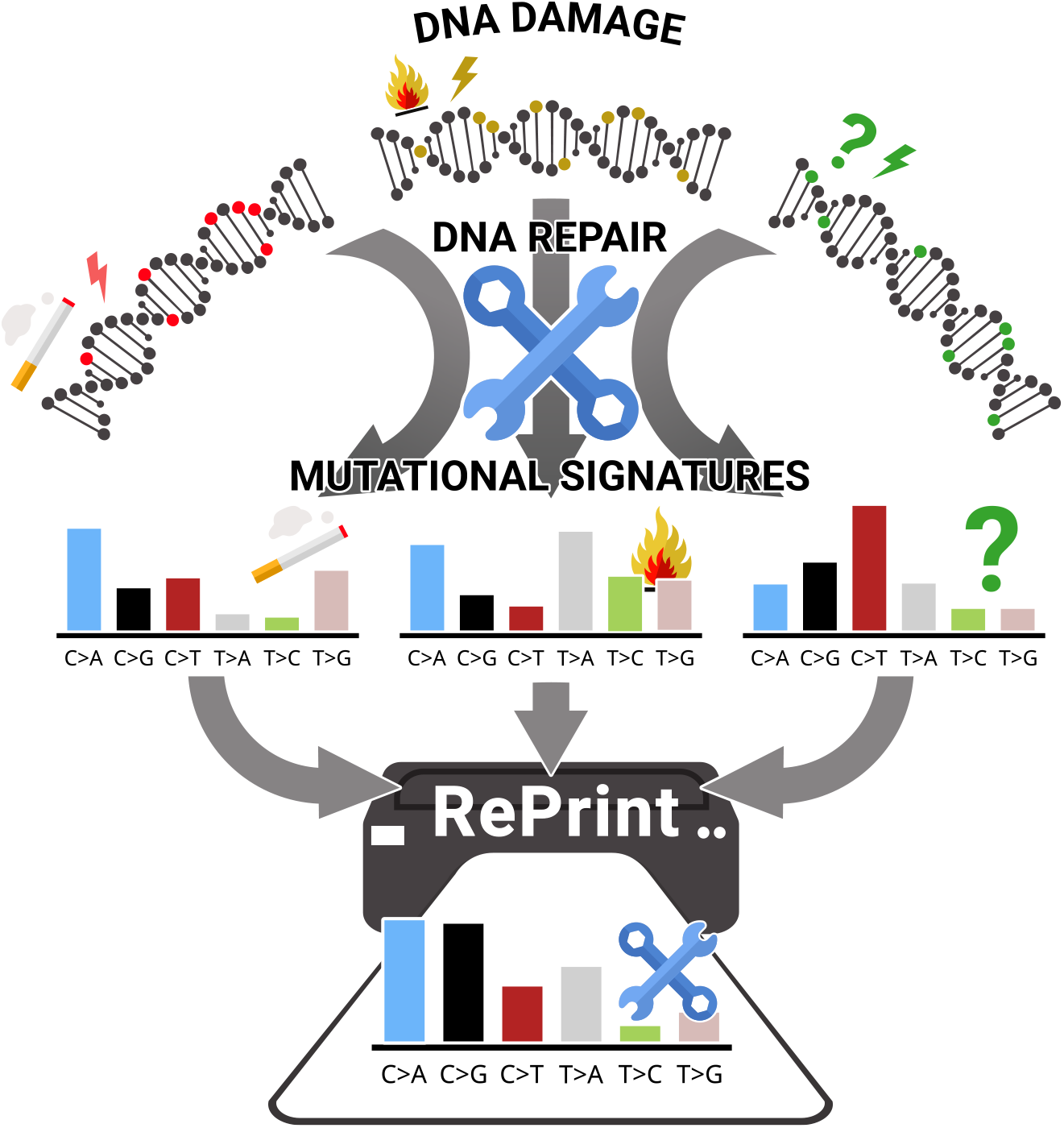
The main idea behind RePrint transformation. Signatures of mutagenic processes are transformed into their RePrints, representing mutation profiles conditioned on the initial damage. Similarity of RePrints suggests related DNA repair mechanisms. In the figure, the first two mutagens with distinct mutational signatures (representing smoking and aromatic hydrocarbons related to incomplete combustion of organic materials) have very similar RePrints reflecting related DNA damage and repair processes. The third mutagen (right) of unknown origin also has a similar RePrint, allowing us to propose the same DNA repair despite of unknown origin.

While DNA-damaging agents are responsible for introducing a specific pattern of damage, the imperfection in the damage repair mechanism (or its “*mis-repair*” tendencies) decides which of these mutagen-introduced lesions are fixed and which become mutations. Signatures may have the same repair mechanism, while differing in their underlying DNA damage step, as different sources of DNA damage can be processed by the same DNA repair pathway. For example, SBS15 and SBS26 are two different Mismatch Repair deficiency signatures, with SBS15 biased towards GC-rich genomic regions and SBS26 biased towards AT-rich genomic regions. Despite their different genomic locations, DNA repair process in both cases is might be the same (Wojtowicz et al., 2020, 2021). Thus very different signatures can share similar RePrints.

To demonstrate the ability of RePrint similarity to elucidate signatures that share a common DNA repair step, we leveraged an expanded collection of mutational signatures, including novel, experimentally-annotated signatures. Based on these annotations, we identified groups of signatures that could be confidently assumed to have the same DNA repair mechanism and used them as a *gold standard* for testing RePrint. Across several metrics, RePrint similarity consistently outperformed signature similarity in the task of identifying common DNA repair pathways. Finally, we applied a “guilt-by-association” approach to predict DNA repair mechanisms for signatures outside the gold-standard set and validated most of these predictions using supporting evidence from the literature Fig. 1. Collectively, these results demonstrate the power of RePrint to capture relationships between DNA repair processes that underlie distinct mutational signatures.

## 2. Results

### 2.1. Method overview

Cells are equipped with multiple DNA repair pathways that specialize in recognizing and repairing different types of DNA damage. The main DNA repair pathways are summarized in Fig. 2 and include mismatch repair (MMR), nucleotide excision repair (NER), base excision repair (BER), homologous recombination (HR), and non-homologous end joining (NHEJ). In addition, translesion synthesis (TLS) is a cellular process that allows DNA replication to bypass DNA lesions using specialized DNA polymerases, including those induced by UV light.

**Figure 2.**
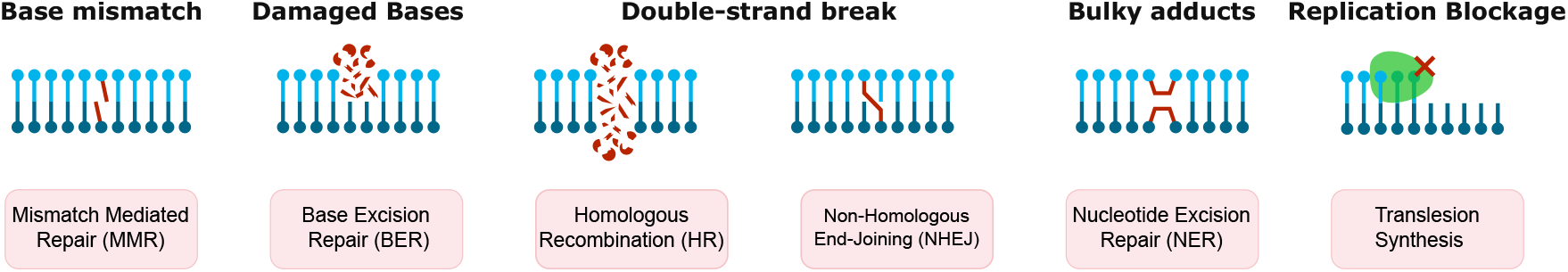
Major DNA repair pathways. Different types of damage (top) are repaired (predominantly) by specific DNA repair pathways (bottom).

Activation of a specific DNA repair pathway is conditional on DNA damage. Therefore, we model the DNA repair step as a transformation from damaged nucleotides due to a mutagenic process to a final mutation pattern after the DNA repair step. We consider the probability that nucleotide *X* mutates to nucleotide *Y*, given two conditions: (1) *X* is in the central position of the trinucleotide context LXR and (2) *X* is known to be mutated. This approach defines the frequency of the three possible mutations for each trinucleotide context, and thus for each context the probabilities sum to one. These conditional probabilities are represented as a vector called RePrint. Formally, RePrint is defined as follows. Let a mutational signature be described as a probability distribution over 96 mutation categories. Each mutation category is determined by a trinucleotide context *L*[*X*→ *Y*]*R*, where:

- *X* ∈ {*C, T* } is the reference base (by convention, purines are converted to their pyrimidine complements on the reverse strand),
- *Y* ∈ {*A, C, G, T* }*\* {*X*} is the variant base,
- *L, R* ∈ {*A, C, G, T*} are the 5’ and 3’ flanking bases, respectively.

Let *S*(*L*[*X* →*Y*]*R*) denote the probability of signature emission, that is, the probability with which the signature generates a mutation category *L*[*X* → *Y*]*N*. The RePrint transforms this signature into a conditional probability distribution. Specifically, for each of the 32 possible trinucleotide contexts *LXR*, RePrint computes the probability of each possible mutation of *X*, given that *X* is mutated:

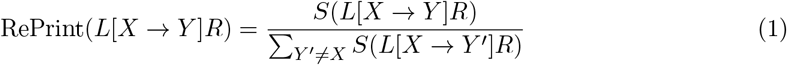

Thus, while a mutational signature is a single categorical probability distribution over 96 mutation types, a RePrint consists of a vector of 32 independent categorical probability distributions: one distribution over 3 possible mutations for each of the 32 trinucleotide contexts. In practice, some trinucleotide contexts may never or rarely be mutated by certain mutagenic processes, which can lead to undefined or noisy RePrint values. To address this, a small pseudocount is added to all signature probabilities before computing the RePrint: *S′*(*L*[*X* → *Y*]*R*) = *S*(*L*[*X* → *Y*]*R*)+*ε*. This ensures that for trinucleotide contexts that are rarely or never mutated in a given signature, the corresponding RePrint conditional probability distribution is approximately uniform, thus preventing over-interpretation of noisy data.

Recall that the RePrint transformation is designed to separate the result of DNA “mis-repair” from the DNA damage component in a mutational signature. By normalizing within each trinucleotide context, RePrint effectively factors out the influence of nucleotide bias in genomic regions that might be preferentially damaged. This can be illustrated by the example of two mutagenic processes that activate the same DNA repair pathway but damage different genomic regions, such as one process preferentially damaging GC-rich regions while another targets AT-rich regions. Their raw mutational signatures would differ substantially because they reflect different distributions of mutated trinucleotides throughout the genome. However, the conditional probabilities captured by the RePrint depend primarily on how the DNA repair machinery processes that particular lesion, not on how frequently that trinucleotide is damaged. Thus, if two signatures share the same DNA repair deficiency but differ in their damage distribution (mutation opportunities), their RePrint will be similar even when their raw signatures are dissimilar. This mathematical property allows the RePrint approach to identify common DNA repair mechanisms underlying diverse mutational processes.

An implementation of the RePrint method is available as an open-source Python package. It computes RePrints directly from input mutational signatures (e.g., COSMIC) while providing a command-line interface for batch processing. It automatically generates visualizations for each signature, produces similarity dendrograms and optional heatmaps, and exports to files. The computed RePrint matrices can also be saved, facilitating downstream analyses and integration with other pipelines.

### 2.2. Evaluation of RePrint similarity as a predictor of common DNA repair pathways

To evaluate the ability of RePrint to identify shared DNA repair mechanisms, we first established a gold standard set of clusters based on precisely annotated signatures. We then used several methods to evaluate how well these clusters are reconstructed using RePrint similarity versus RSMD similarity. Finally, we tested the potential of using the guilt-by-association principle to predict the DNA repair pathway for the signatures not in the gold standard set.

#### 2.2.1. Construction of the gold standard cluster set

To evaluate RePrint’s ability to identify shared DNA repair mechanisms across signatures derived from diverse mutagenic processes, we used mutational signatures from three sources. We included the COSMIC version 3.4 signature set (human genome GRCh37 as the primary data set. This set was extended with environmental mutational signatures from Kucab et al. (2019), who systematically characterized signatures arising from exposure to various environmental agents, including polycyclic aromatic hydrocarbons (PAHs), nitro-PAHs, aristolochic acid, and other carcinogens using controlled *in vitro* experiments. We further extended the signature set with experimentally-derived signatures from Zou et al. (2021), who used CRISPR-Cas9-based knockouts of DNA repair genes in isogenic cell models to generate signatures associated with specific repair pathway deficiencies, including knockouts of MMR genes (MSH6, PMS1, PMS2), HR-related genes (EXO1, RNF168), and other DNA repair components. All mutational signatures from the additional data sets are listed in Supplementary Tables S1 and S2.

We utilized validated COSMIC signatures and signatures experimentally induced by related environmental mutagens (Kucab et al., 2019) or by disruption of genes within the same DNA repair pathway (Zou et al., 2021) to construct a two-tier hierarchy of reference signature clusters. *Homogeneous clusters* comprise signatures that are known to arise from the same or highly similar DNA damage sources, either chemically (e.g., PAHs, ROS) or mechanistically (e.g., MMRd, HRD), and are expected to engage similar DNA repair pathways. *Broader clusters* combine multiple homogeneous clusters based on a shared DNA repair pathway, while allowing for more diverse underlying mutagenic processes. This hierarchical structure enables us to test whether RePrint can identify common repair mechanisms not only among highly similar signatures but also among more divergent signatures that share repair machinery but differ in their signature profiles. The clusters are summarized in Table 1.

**Table 1:** Annotated reference (gold standard) signature clusters. *Homogeneous Clusters* are defined as sets of signatures known to originate from the same or very similar DNA damage and activate a similar repair pathway. Some signatures in this cluster are experimentally induced by similar environmental mutagens (Kucab et al., 2019) or disruption of the same DNA repair pathway (Zou et al., 2021). *Broad Clusters* of signatures contain signatures that are assumed to be mainly repaired by a common DNA repair pathway, but allowing for more heterogeneous DNA damage processes. Lists of all environmental signatures and gene knockout signatures are in Supplementary Tables S1 and S2, respectively.

| Cluster | Signatures | Description |
| --- | --- | --- |
| **Homogeneous Clusters** |  |  |
| PAHs | SBS4, 8 environmental signatures | Polycyclic aromatic hydrocarbons produced by incomplete combustion of organic matter, including tobacco. Bulky adducts repaired by NER. |
| NitroPAHs | 4 environmental signatures | Nitrated PAHs produced by oxidation of PAHs by nitrate radicals ($\text{NO}_3$). Like PAHs, NitroPAHs form bulky DNA adducts. |
| ROS | SBS18, SBS36, SBS38 | Reactive Oxygen Species: SBS18 and SBS36 are directly linked to ROS; SBS38 is associated with indirect UV exposure generating ROS. |
| HRD | SBS3 and two signatures of HR-related gene knockouts | Homologous recombination deficiency: SBS3 and $\Delta\text{EXO1}$, $\Delta\text{RNF168}$ mutational signatures. |
| TMZ | SBS11 and two TMZ environmental signatures | Temozolomide treatment. |
| Platinum | SBS31, SBS35, and two environmental signatures | Platinum drug treatment. |
| AAs | SBS22a, SBS22b, and one environmental signature | Aristolochic Acid treatment. |
| MMRd | SBS6, SBS14, SBS15, SBS20, SBS21, SBS26, SBS44, and three MMR gene knockout signatures | DNA Mismatch Repair deficiency: $\Delta\text{MSH6}$, $\Delta\text{PMS1}$, $\Delta\text{PMS2}$ mutational signatures. |
| **Broad Clusters** |  |  |
| PAHs<sub>broad</sub> | PAHs, NitroPAHs | Air pollutants from incomplete combustion of polycyclic and nitrated PAHs. |
| MMRd<sub>broad</sub> | MMRd, Temozolomide | Direct or indirect MMRd; includes Temozolomide due to evidence that it alters mismatch repair and increases mutational burden. |
| DrugsBA<sub>broad</sub> | Platinum, AA, SBS24 | DNA damage from drug treatments producing bulky DNA adducts. |

#### 2.2.2. Evaluation of RePrint-based hierarchical clustering of signatures using gold standard clusters

To assess how well RePrint similarity captures shared DNA repair mechanisms compared to signature similarity, we performed hierarchical clustering analysis on both the similarity of signatures and similarity of their corresponding RePrints (Fig. 3). For each representation, we constructed a distance matrix using root-mean-square deviation (RMSD) between all pairs, then applied unsupervised hierarchical clustering with complete linkage and Euclidean distance. We evaluated the resulting dendrograms using three complementary quantitative metrics: (1) Dasgupta’s objective score, (2) an intra-cluster compactness score, and (3) an inter-cluster separation score. These metrics allowed us to systematically compare clustering quality between signature-based and RePrint-based representations while accounting for the known biological relationships encoded in our gold standard reference clusters.

**Figure 3.**
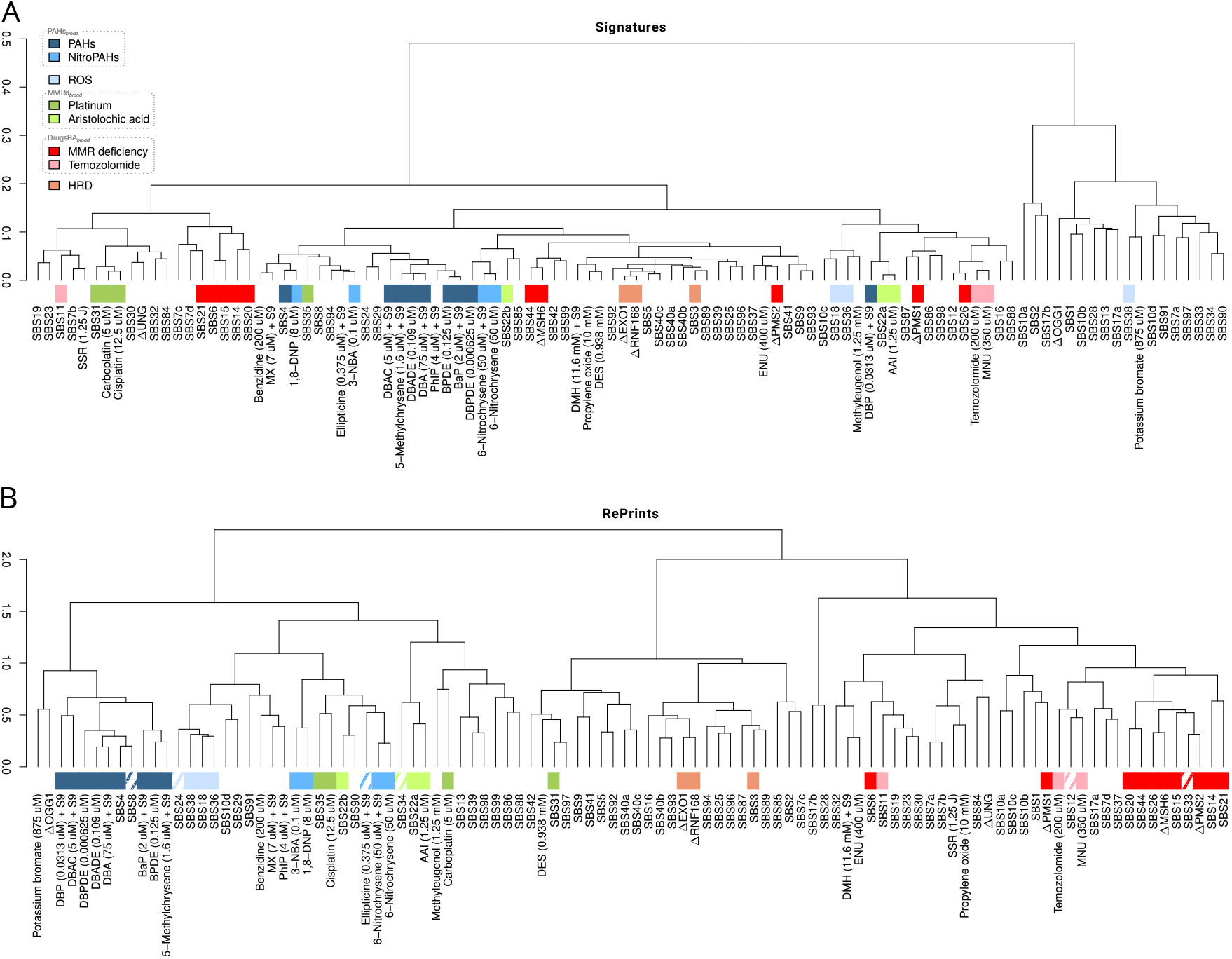
RePrint clustering captures shared DNA repair mechanisms. Hierarchical clustering dendrograms of (A) mutational signatures and (B) RePrints for COSMIC, environmental exposure, and gene knockout signatures. Colored boxes indicate gold standard (homogeneous and broad) cluster annotations (see legend, panel A), while hatched boxes show *guilt-by-association* predictions. RePrint-based clustering (B) more effectively groups signatures by shared repair mechanisms than standard signature clustering (A), enabling the inference of DNA repair pathways for signatures of unknown origin.

Dasgupta’s objective score (Dasgupta, 2016) provides a way to evaluate hierarchical clustering quality by penalizing the placement of dissimilar items near each other in the tree (and thus is independent of a gold standard clusters). For each pair of items, the score accumulates a penalty equal to their dissimilarity weighted by the size of the smallest subtree containing both items (i.e., the size of their lowest common ancestor’s subtree). We define our dissimilarity function to reflect the hierarchical structure of our gold standard clusters: pairs from the same homogeneous cluster receive a dissimilarity value of 0, pairs from the same broader cluster but different homogeneous clusters receive a dissimilarity value of 1, and pairs from different broader clusters receive a dissimilarity value of 2. Under this scoring scheme, a good hierarchical clustering should yield a low total score, indicating that similar signatures (by our gold standard) cluster together at low hierarchical levels while dissimilar signatures separate at higher levels. We compute Dasgupta’s objective for both the signature-based and RePrint-based clustering dendrograms and compare the results across all reference clusters (Figure 4).

**Figure 4.**
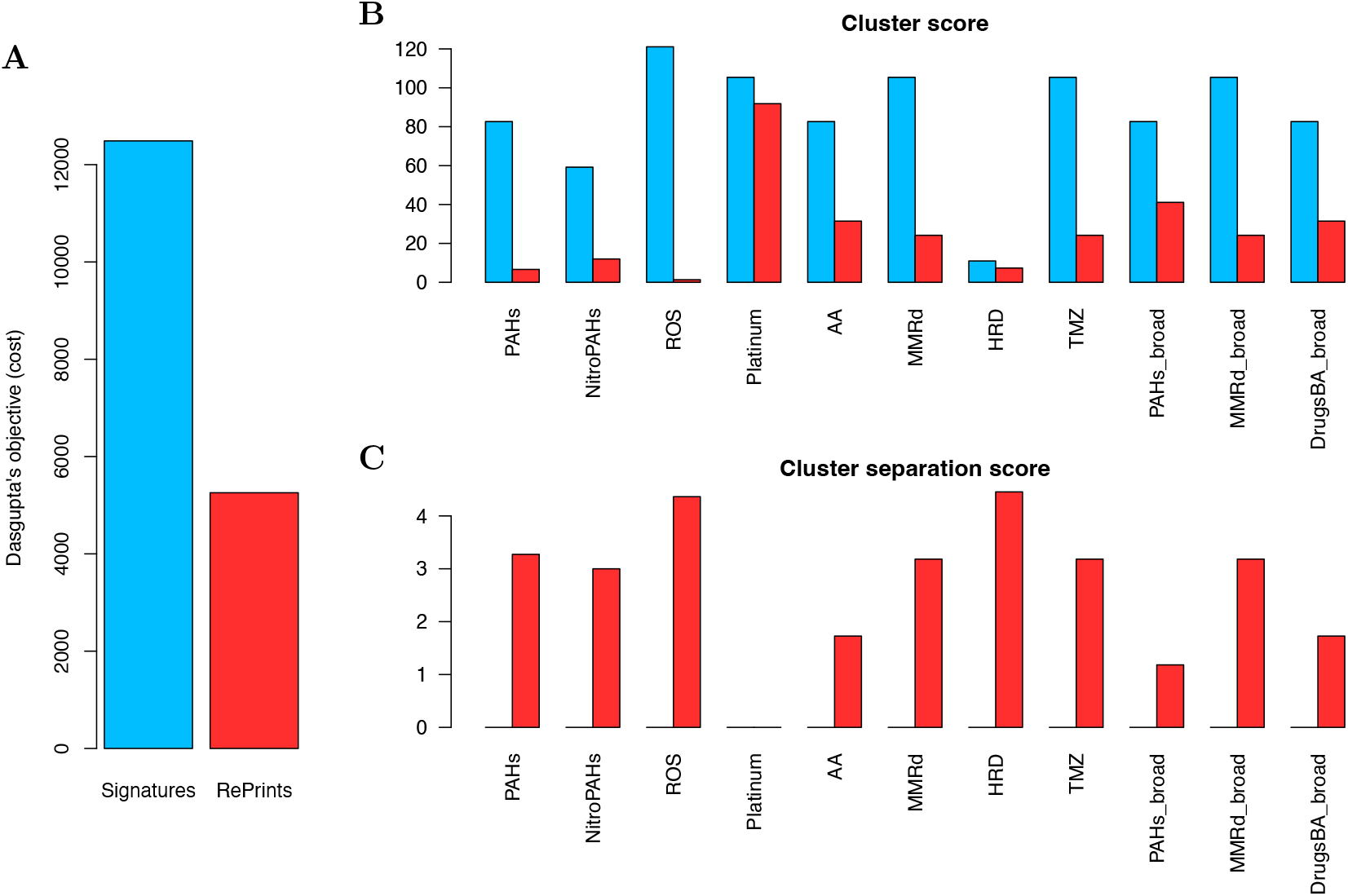
Evaluation of gold standard cluster recovery. RePrint-based representation (red) is compared against mutational signature-based representation (blue), both clustered using hierarchical clustering with complete linkage and Euclidean distance on RMSD distance matrices. (A) Dasgupta’s objective measuring overall dendrogram quality (higher is better). (B) Intra-cluster compactness across individual gold standard clusters (lower is better). (C) Inter-cluster separation score (higher is better).

To quantify how tightly members of each reference cluster group together in hierarchical clustering, we computed an intra-cluster compactness score using `pvclust` (Suzuki and Shimodaira, 2006), which calculates p-values for hierarchical clustering via multiscale bootstrap resampling (see details in Supplementary Figure S2). These p-values represent the statistical confidence that each cluster represents a true grouping rather than arising by chance. For each of our reference clusters (both homogeneous and broad), we identified the smallest subtree containing all of its members, which corresponds to their lowest common ancestor (LCA), and computed the intra-cluster compactness score as the sum of negative log p-values over internal branches of this subtree. Here, smaller scores indicate stronger support for the grouping of cluster members. This metric thus captures whether signatures expected to share DNA repair mechanisms cluster together with high statistical confidence, providing a measure of cluster cohesiveness that accounts for sampling variability.

To complement the intra-cluster compactness measure, we computed an inter-cluster separation score that quantifies how well different reference clusters are distinguished in the hierarchical tree. For each pair of reference clusters *C*_*i*_ and *C*_*j*_, we identified their respective LCA nodes in the dendrogram. We then calculated the separation between these clusters as the number of edges along the path in the tree connecting their two LCA nodes. This measure captures the hierarchical distance at which two clusters diverge: pairs of clusters that separate early in the tree (near the root) receive higher separation scores, while pairs that remain together until lower levels receive lower scores. The overall inter-cluster separation score for a given reference cluster is computed by averaging its separation values from all other reference clusters. Higher separation scores indicate that the reference cluster is well-distinguished from other clusters in the hierarchy, suggesting that the clustering successfully captures distinct biological groupings.

The results of all scoring functions are summarized in Fig. 4. RePrint outperformed signature similarity for all tests supporting the hypothesis that it provides a better measure of similarity of DNA repair pathways than signature similarity.

### 2.3. Prediction of DNA repair pathway using guilt-by-association principle

In order to predict DNA repair pathways for signatures not explicitly assigned in the reference set, we utilize a guilt-by-association approach. Specifically, we apply guilt-by-association when the query signature is the only unknown signature in a subtree of at least 3 signatures, where all remaining signatures are from the same gold standard cluster. Based on this strategy, we predicted the repair mechanism of mutational signatures SBS8, SBS24, SBS90, SBS33, SBS12, and SBS34, as well as of the Ellipticine signature. A summary of the pathway predictions along with corresponding supporting evidence can be found in Table 2. Visualizations of RePrints for examples of signatures with predicted with the guilt-by-association principle, along with their closest RePrint neighbours, are shown in Fig. 5. Five our of six predictions were confirmed based on literature, and the correctness of the sixth could not be confirmed or rejected.

**Table 2:** Summary of predictions and supporting evidence for signatures and compounds.

| Query signature | Associated annotated cluster | Proposed DNA damage/repair pathway | Validation | Comments on the validation |
| --- | --- | --- | --- | --- |
| SBS8 | PAHs | DNA adducts/NER | Jager et al. (2019) Holcomb et al. (2016) | SBS8 can be obtained by Ercc1 (NER pathway gene) knockout; PAHs create bulky adducts and tobacco smoking inhibits NER |
| SBS24 | ROS | ROS | Huang et al. (2020) | Aflatoxin induces oxidative stress by increasing ROS. |
| Ellipticine | NitroPAHs | DNA adducts/NER | Stiborova et al. (2004) | Anticancer drug involving DNA adduct formation |
| SBS33 | MMRd/MMRd<sub>broad</sub> | MMRd | Levatić et al. (2022) | Proposes a relation between SBS33 and MMRd |
| SBS12 | MMRd/MMRd<sub>broad</sub> | MMRd | Mas-Ponte et al. (2022) | SBS12 can be obtained by PMS2 (MMRd gene) knockout |
| SBS34 | DrugsBA<sub>broad</sub> | DNA adducts/NER | None | Signature of unknown origin. |

**Figure 5.**
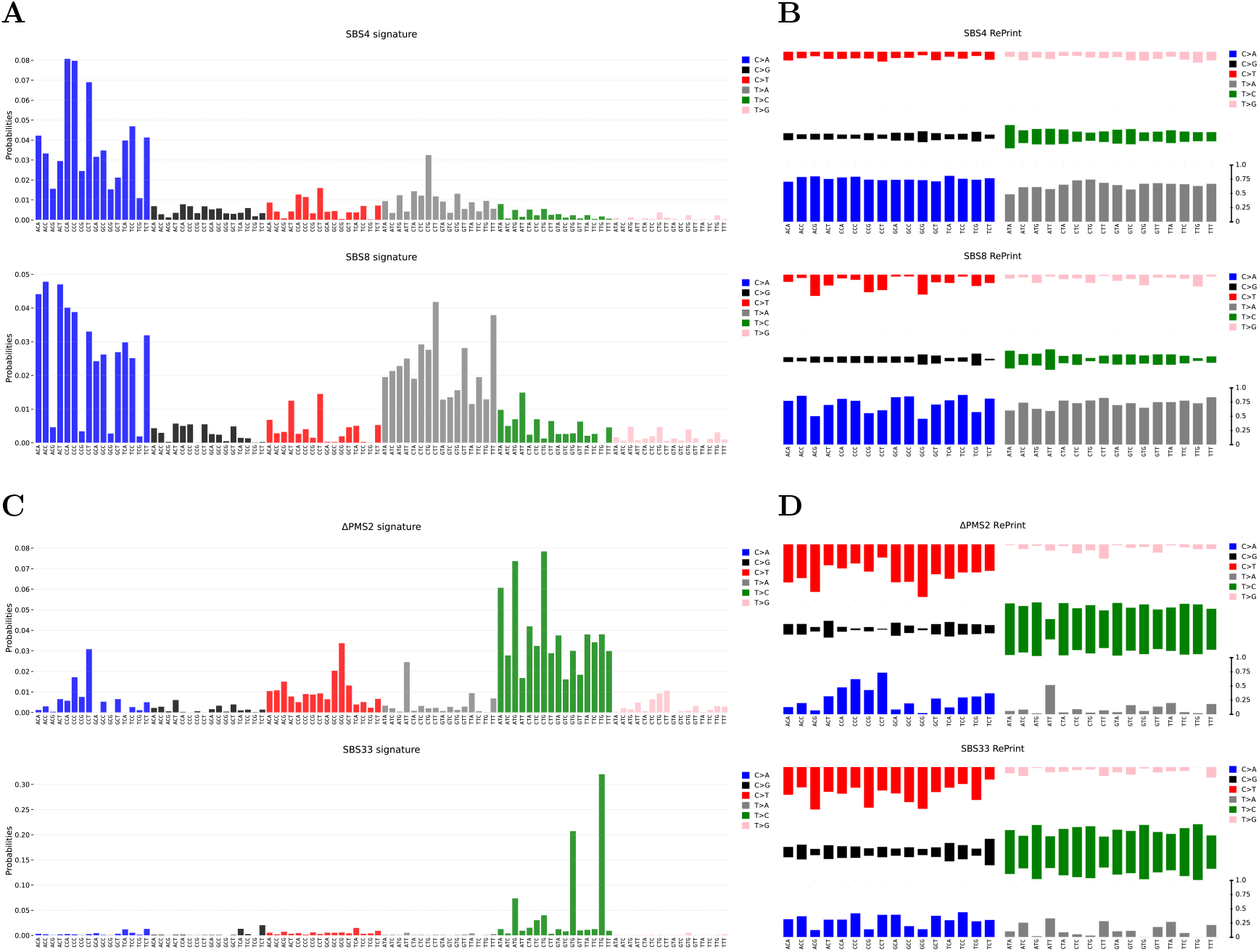
Mutational signatures and RePrint visualizations. Left panels (A,C) show mutational signatures as probability distributions over 96 mutation categories. Right panels (B,D) show the corresponding RePrints, which transform signatures into conditional probabilities (bars sum to one within each trinucleotide context). RePrint similarity revealed that SBS8 clusters with SBS4 (B) and SBS33 clusters with ΔPMS2 knockout (D) despite different mutational profiles (A,B), suggesting their shared DNA repair pathways within each pair through *guilt-by-association*.

In contrast to RePrint-based guilt-by-association, the use of signature RSMD -similarity-based guilt-by-association with the same clustering criterion allowed for the prediction of repair pathways for only two signatures, one of which (SBS12) was also predicted by RePrint. The second (PhIP – Heterocyclic amines (HCAs)) clustered with PAHs and belongs to a group of mutagenic compounds produced during thermal processing of protein-rich foods Zhao et al. (2021). It is chemically different from PAHs, binds DNA differently than PAHs Broyde et al. (2007), but similarly to PAHs, it forms bulky adducts. Thus while the DNA repair pathway for PhIP might not be exactly the same as for PAHs, it is generally related to DNA adducts repair. This suggests that RSMD similarity is also informative but less powerful than RePrint.

## 3. Discussion

Understanding the mechanisms of DNA repair following exposure to mutagens is fundamental for the development of effective therapies. Consequently, the ability to infer DNA repair pathways involved in repairing damage caused by mutagenic processes in cancer genomes is of primary importance. However, the etiology of many mutational signatures remains unclear, making it challenging to identify the underlying DNA repair mechanisms. RePrint fills this gap by enabling the inference of such mechanisms using the principle of guilt-by-association.

Analysis of RePrint clusters also yields several noteworthy insights. For example, consistent with previous findings (Wojtowicz et al., 2020), RePrints of MMRd signatures cluster together. This is consistent with the notion that Mismatch Repair Deficiency generally activates the same DNA repair pathway independently of the specific MMRd signature. Furthermore, co-clustering of MMRd signatures with signatures associated with Temozolomide (TMZ) treatment reinforces the idea that RePrints effectively capture underlying DNA repair processes since tumors that are responsive to TMZ have been proposed to induce MMRd (Crisafulli et al., 2022). In addition, signatures (SBS10a,b) are in this group which is explained by the fact that errors associated with polymerase epsilon deficiency are assumed to be primarily repaired through mismatch repair (MMR) Andrianova et al. (2017) and thus the pattern of un-repaired lesions is expected to be consistent with MMRd. In contrast, only one of the two signatures related to polymerase delta (SBS10c) is contained in this cluster, suggesting that SBS10d, which is associated with defective POLD1 proofreading, relies on a different repair pathway.

As another example, polycyclic aromatic hydrocarbons (PAHs), are a class of environmental pollutants that lead to the formation of DNA adducts. The main pathway involved in repairing this DNA damage is NER. As a predominant component of cigarette smoke, the co-clustering of SBS4 with PAHs confirms the power of RePrint to capture shared DNA repair pathways. Importantly, SBS8, a signature that is still annotated to be of unknown origin in COSMIC, co-clusters with PAHs, suggesting its relation to the NER pathway. This observation is consistent with two previous studies linking SBS8 to a deficiency in the NER pathway: The first provided experimental evidence that deletion of the ERRC2 gene (a key component of the NER pathway) generates a signature similar to SBS8 (Jager et al., 2019), while the second study shows that SBS8 is associated with down-regulation of several NER genes (Amgalan et al., 2023).

As a final example, reactive oxygen species (ROS) are generated in cells as byproducts of normal endogenous metabolic processes, UV radiation, and exogenous chemicals (Cooke et al., 2003; de Jager et al., 2017). As expected, RePrints of known ROS-related signatures clustered together. Our ROS cluster is seeded by three signatures: SBS18, SBS38, and SBS36. SBS18 and SBS36 are the two mutational signatures attributed directly to reactive oxygen species (Pilati et al., 2017; Viel et al., 2017) while SBS38 is associated with indirect ultra-violet exposure and thus subject to ROS generation by UV light (de Jager et al., 2017). These three signatures are co-clustering with SBS24 - a signature associated with Aflatoxin exposure. Aflatoxins are mutagens that are produced by certain molds and are proposed to induce oxidative stress by increasing reactive oxygen species (ROS) (Huang et al., 2020). The co-clustering of SBS24 with ROS-related signatures supports this view.

Our graphical visualization of RePrints (Fig. 5) often exposes striking properties of signatures. For example, it can be observed that in SBS4 about 60% cases when C or T is mutated it is mutated to A independently of the trinucleotide context (Fig. 5). Such observations are likely to contribute to a better understanding of DNA repair process associated with a given signature.

Although the analysis presented in this study highlights the strength of RePrint, it should be noted that the approach has certain limitations. In particular, for signatures in which some triples are never (or rarely) mutated, the respective conditional probability of RePrint might not be sufficiently informative, leading to potential false positive and false negative results. However, these notwithstanding, RePrint provides the first approach to transfer information between signatures, opening the door to inferring the properties of signatures of unknown origin and providing a promising tool for machine learning applications.

## Acknowledgment

This research was supported in part by the Division of Intramural Research of the National Library of Medicine (NLM), National Institutes of Health (NIH). The contributions of the NIH authors (J.H., T.M.P.) are considered Works of the United States Government. The findings and conclusions presented in this paper are those of the authors and do not necessarily reflect the views of the NIH or the U.S. Department of Health and Human Services. This work was also supported by the Polish National Agency for Academic Exchange via Polish Returns Program (D.W.) and the US-Israel Binational Science Foundation (R.S.).

