## Supplementary Information for "Toward identification of common DNA repair process in mutational signatures"

Table S1: List of 53 mutagen-induced substitution signatures (with treatment conditions) studied in (Kucab et al., 2019). We removed some signatures from our analysis—if a signature was not stable (usually due to a low number of mutations) or if it was similar to its variant at a different concentration (see Figure 3B in (Kucab et al., 2019)). The last column indicates with which homogeneous cluster a signature was annotated with.

| Mutagen/Treatment | Group | Comment | Homogeneous cluster |
|---|---|---|---|
| Formaldehyde (120 $\mu$M) | Aldehydes | Excluded. Unstable signature due to low mutation count. | |
| MNU (350 $\mu$M) | Alkylating agents | | TMZ |
| ENU (400 $\mu$M) | Alkylating agents | | |
| DMS (0.078 mM) | Alkylating agents | Excluded. Unstable signature due to a low mutation count. | |
| DES (0.938 mM) | Alkylating agents | | |
| DMH (11.6 mM) + S9 | Alkylating agents | | |
| Temozolomide (200 $\mu$M) | Drug therapy | | TMZ |
| Temozolomide (200 $\mu$M) | Drug therapy | Excluded. A replicate of included signature. | |
| Cyclophosphamide (18.75 $\mu$M) + S9 | Drug therapy | Excluded. Unstable signature due to a low mutation count. | |
| Mechlorethamine (0.3 $\mu$M) | Drug therapy | Excluded. Unstable signature due to a low mutation count. | |
| Semustine (150 $\mu$M) | Drug therapy | Excluded. Unstable signature due to a low mutation count. | |
| Cisplatin (3.125 $\mu$M) | Drug therapy | Excluded. Replicate at a different concentration is included. | |
| Cisplatin (12.5 $\mu$M) | Drug therapy | | Platinum |
| Carboplatin (5 $\mu$M) | Drug therapy | | Platinum |
| Ellipticine (0.375 $\mu$M) + S9 | Drug therapy | | |
| AZD7762 (1.625 $\mu$M) | DNA damage response inhibitor | Excluded. Unstable signature due to a low mutation count. | |
| Benzidine (200 $\mu$M) | Aromatic amines | | |
| 4-ABP (300 $\mu$M) + S9 | Aromatic amines | Excluded. Unstable signature due to a low mutation count. | |
| PhIP (3 $\mu$M) + S9 | Heterocyclic amines | Excluded. Replicated at a different concentration is included. | |
| PhIP (4 $\mu$M) + S9 | Heterocyclic amines | | |
| SSR (1.25 J) | Radiation | | |
| N-Nitrosopyrrolidine (50 mM) | Nitrosamine | Excluded. Unstable signature due to a low mutation count. | |

Continued on next page

Table S1 – continued from previous page

| Mutagen/Treatment | Group | Comment | Homogeneous cluster |
|---|---|---|---|
| BaP (0.39 $\mu$M) + S9 | PAHs | Excluded. Replicate at a different concentration is included. | |
| BaP (2 $\mu$M) + S9 | PAHs | | PAHs |
| BPDE (0.125 $\mu$M) | PAHs | | PAHs |
| DBA (75 $\mu$M) + S9 | PAHs | | PAHs |
| DBADE (0.0313 $\mu$M) | PAHs | Excluded. Replicate at a different concentration is included. | |
| DBADE (0.109 $\mu$M) | PAHs | | PAHs |
| DBP (0.0039 $\mu$M) | PAHs | Excluded. Unstable signature due to a low mutation count. | |
| DBP (0.0313 $\mu$M) + S9 | PAHs | | PAHs |
| DBPDE (0.000156 $\mu$M) | PAHs | Excluded. Replicate at a different concentration is included. | |
| DBPDE (0.000625 $\mu$M) | PAHs | | PAHs |
| 5-Methylchrysene (1.6 $\mu$M) + S9 | PAHs | | PAHs |
| DBAC (5 $\mu$M) + S9 | PAHs | | PAHs |
| 6-Nitrochrysene (0.78 $\mu$M) | Nitro-PAHs | Excluded. Replicate at a different concentration is included. | |
| 6-Nitrochrysene (50 $\mu$M) | Nitro-PAHs | | NitroPAHs |
| 6-Nitrochrysene (12.5 $\mu$M) + S9 | Nitro-PAHs | Excluded. Replicated at a different concentration is included. | |
| 6-Nitrochrysene (50 $\mu$M) + S9 | Nitro-PAHs | | NitroPAHs |
| 1,6-DNP (0.09 $\mu$M) | Nitro-PAHs | Excluded. Similar to 1,8-DNP. | |
| 1,8-DNP (0.125 $\mu$M) | Nitro-PAHs | Excluded. Replicated at a different concentration is included. | |
| 1,8-DNP (8 $\mu$M) | Nitro-PAHs | | NitroPAHs |
| 3-NBA (0.025 $\mu$M) | Nitro-PAHs | Excluded. Unstable signature due to a low mutation count. | |
| 3-NBA (0.1 $\mu$M) | Nitro-PAHs | | NitroPAHs |
| Potassium bromate (260 $\mu$M) | ROS | Excluded. Replicated at a different concentration is included. | |
| Potassium bromate (875 $\mu$M) | ROS | | |
| AAI (1.25 $\mu$M) | others | | AAs |
| AAII (37.5 $\mu$M) | others | Excluded. Unstable signature due to a low mutation count. | |
| AFB1 (0.25 $\mu$M) + S9 | others | Excluded. Unstable signature due to a low mutation count. | |
| OTA (0.08 $\mu$M) + S9 | others | Excluded. Unstable signature due to a low mutation count. | |
| Propylene oxide (10 mM) | others | | |
| Furan (100 mM) + S9 | others | Excluded. Unstable signature due to a low mutation count. | |
| Methyleugenol (1.25 mM) | others | | |
| MX (7 $\mu$M) + S9 | others | | |

Continued on next page

Table S1 – continued from previous page

| Mutagen/Treatment | Group | Comment | Homogeneous cluster |
|---|---|---|---|

Table S2: List of 9 isogenic CRISPR–Cas9 gene knockouts from Zou et al. (2021) that produced marked mutational signatures. If a signature was very similar to that of a different knockout, then it was not included in our analysis (see Figure 3a in (Zou et al., 2021)). The last column indicates with which homogeneous cluster a signature was annotated.

| Gene knockout | Pathway | Comment | Homogeneous cluster |
|---|---|---|---|
| EXO1 | HR | | HRD |
| MLH1 | MMR | Not included due to its similarity to ΔMSH6 signature. | |
| MSH2 | MMR | Not included due to its similarity to ΔMSH6 signature. | |
| MSH6 | MMR | | MMRd |
| OGG1 | BER | | |
| PMS1 | MMR | | MMRd |
| PMS2 | MMR | | MMRd |
| RNF168 | DSB repair & Checkpoint | | HRD |
| UNG | BER | | |

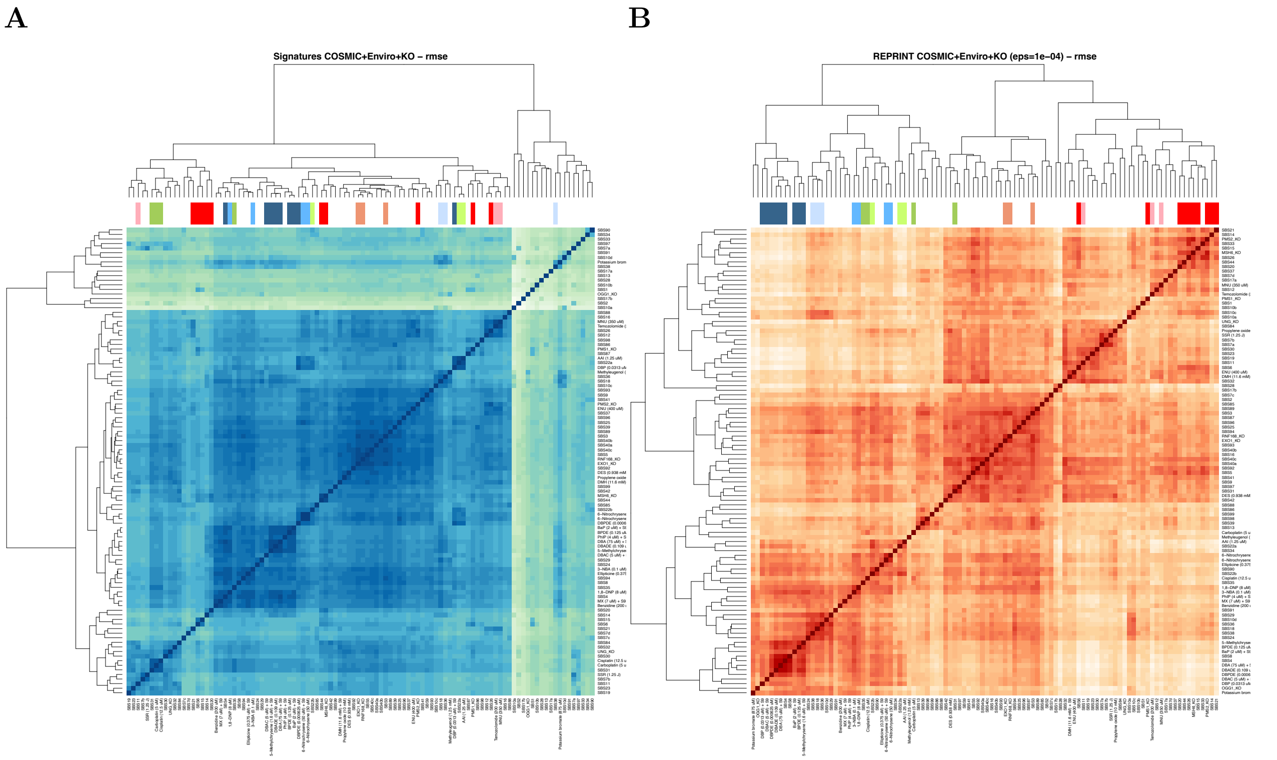


Figure S1: **Comparison of mutational signature and RePrint similarity heatmaps.** Hierarchical clustering heatmaps of pairwise RMSD distances between (A) mutational signatures and (B) RePrints for COSMIC, environmental exposure, and gene knockout signatures. Darker colors indicate higher similarity. Colored bars denote reference gold standard cluster annotations (see Figure 3 for explanations).

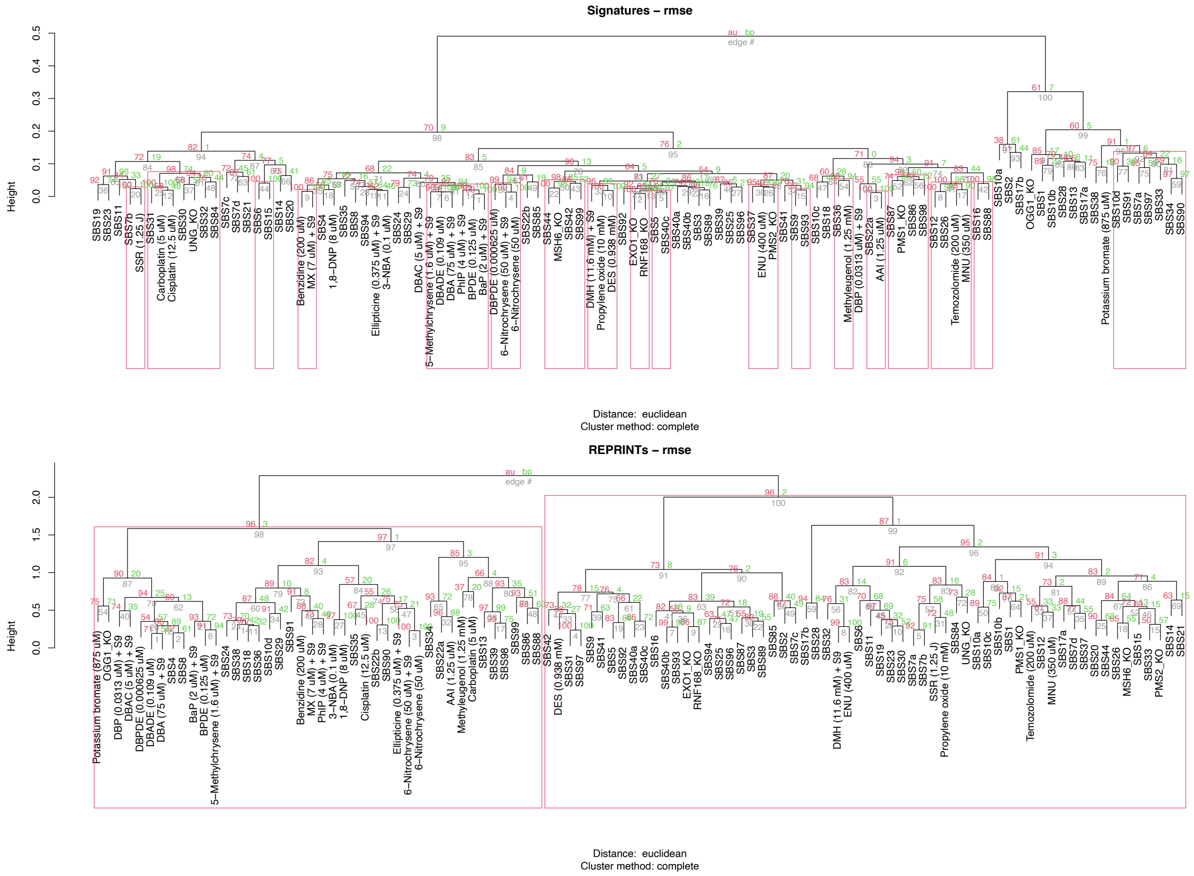


Figure S2: **P-values for dendrograms in Figure 3.** For similarity of both mutational signatures (top) and RePrint (bottom), `pvclust` performed hierarchical clustering, then computed p-values (approximately unbiased (AU) p-value) for each cluster in the resulting dendrogram through 1000 bootstrap replicates. These p-values represent the statistical confidence supported by the data that each cluster represents a true grouping rather than arising by chance.
